# Simultaneous measurement of nascent transcriptome and translatome using 4-thiouridine metabolic RNA labeling and translating ribosome affinity purification

**DOI:** 10.1101/2023.02.15.525786

**Authors:** Hirotatsu Imai, Daisuke Utsumi, Hidetsugu Torihara, Kenzo Takahashi, Hidehito Kuroyanagi, Akio Yamashita

**Author notes:** Corresponding authors: Hirotatsu Imai, Akio Yamashita.

## Abstract

Regulation of gene expression in response to various biological processes, including extracellular stimulation and environmental adaptation requires *de novo* RNA synthesis and translation. Analysis of the coordinated and circular regulation of dynamic RNA synthesis and translation is required to determine functional protein production. However, reliable methods for the simultaneous measurement of nascent RNA synthesis and translation at the gene level are limited. Here, we developed a novel method for the simultaneous assessment of nascent RNA synthesis and translation by combining 4-thiouridine (4sU) metabolic RNA labeling and translating ribosome affinity purification (TRAP) using a monoclonal antibody against evolutionarily conserved ribosomal P-stalk proteins. The P-stalk-mediated TRAP (P-TRAP) technique recovered endogenous translating ribosomes, allowing easy translatome analysis of various eukaryotes. We validated this method in mammalian cells by demonstrating that acute unfolded protein response (UPR) in the endoplasmic reticulum (ER) induces dynamic reprograming of nascent RNA synthesis and translation. Our method can serve as a simple and powerful tool for analyzing the coordinated regulation of transcription and translation of individual genes in various eukaryotes.

## Introduction

The regulation of gene expression plays a pivotal role in diverse biological and physiological processes such as cell differentiation, development, environmental responses, and immune responses. One determinant of gene expression is the amount of mature RNA, which reflects the balance between RNA synthesis and degradation (1-5). To understand the dynamics of RNA synthesis and degradation comprehensively, metabolic RNA labeling techniques using nucleotide analogs have been developed (6-14). For example, thiol(SH)-linked alkylation for the metabolic sequencing of RNA (SLAMseq) enables to uncover 4-thiouridine (4sU)-labeled transcripts in cDNA libraries by bioinformatic detection of specific T-to-C (T>C) conversions at the sites of 4sU incorporation (9). Combining SLAMseq with QuantSeq, which is a deep sequencing close to the 3’ end of polyadenylated RNAs (15), allows rapid and quantitative access to 4sU-labeled transcripts expression profiles (9). These techniques have addressed the dynamic aspects of gene expression regulation in various biological processes such as RNA quality control, oncogenesis, and embryogenesis (16-18).

The final output of gene expression, which indicates the expression level of proteins, is significantly influenced by RNA translation. Several studies have reported that the primary set of RNA transcripts obtained by standard RNA-seq experiments may have a low quantitative correlation with the proteome (4,19-21). To overcome this issue, analyzing the translatome, which refers to mature RNAs bound to ribosomes for protein synthesis, helps in estimating individual gene expression levels (22-24). To analyze the translatome, ribosome-bound RNAs were fractionated using sucrose density gradients and analyzed using RNA-seq (25,26). Translating ribosome affinity purification (TRAP) was also used to analyze translatomes which requires the expression of affinity-tagged ribosomal proteins of the large (60S) ribosomal subunit by genetic modification of cells and organisms. The cell type-specific promoter controls the expression of affinity-tagged ribosomal proteins, allowing the capture of RNAs bound to ribosomes from specific cells/tissues in organisms (22,27-29). Following the immunoprecipitation of affinity-tagged ribosomes, ribosome-bound RNA was analyzed using RNA-seq. In a recently established ribosome profiling (Ribo-seq) technique, RNase digestion of ribosome-bound RNAs resulted in ribosome-protected RNA fragments (RPFs). The RPFs were converted to a library for deep sequencing to determine the precise position and density of ribosomes at nucleotide resolution (30-32).

Although RNA synthesis, degradation, and translation control are distinct steps, these regulations interplay in gene expression (33,34). The simultaneous measurement of these different hierarchical controls takes advantage of a comprehensive understanding of complicated gene expression regulation. Recently, nascent Ribo-seq (nRibo-seq) was developed to simultaneously measure nascent RNA synthesis and translation by combining Ribo-seq with 4sU metabolic RNA labeling (35). nRibo-seq estimates whether a short RPF is derived from a 4sU-labeled nascent RNA by a binomial distribution and simultaneously measures the RNA synthesis rate and translation efficiency at the level of bulk or specific groups of RNA. This pioneering technique revealed variable ribosome loading rates among the functional gene subsets. However, Ribo-seq deals with the short length of RPFs, making the reliable quantification of 4sU incorporation for individual genes difficult (35).

In this study, we developed a simple method for the simultaneous measurement of nascent polyadenylated RNAs and their translation at the gene level by combining the TRAP technique with 4sU metabolic RNA labeling. Since TRAP captures full-length RNAs bound to the ribosomes, it is compatible with QuantSeq, which analyzes the inherently uridine-rich 3’ UTRs of polyadenylated RNAs, facilitating the robust quantification of T>C conversions. In contrast to the conventional TRAP technique that requires genetically engineered cells, we developed an endogenous ribosome affinity purification method using a monoclonal antibody against an evolutionarily conserved ribosomal P-stalk protein. The P-stalk mediated translational ribosome affinity purification (P-TRAP) technique enables convenient access to the translatome in diverse eukaryotes, including mammals, fish, insects, and nematodes, without genetic manipulations. We applied a combination of conventional TRAP or the newly developed P-TRAP with 4sU metabolic RNA labeling and defined the nascent transcripts and their translation at the level of individual genes in the acute unfolded protein response (UPR) within the endoplasmic reticulum (ER). This simple and versatile translatome analysis technique would enhance our understanding of the complex regulation of gene expression in various eukaryotes.

## Materials and Methods

### Plasmid and antibodies

A plasmid for the stable expression of enhanced green fluorescent protein (EGFP) fused to the N-terminus of the ribosomal protein L10a (EGFP-L10a) in mammalian cells was constructed using standard procedures. Briefly, a synthetic DNA fragment of EGFP-L10a was cloned into a expression vector pcDNA5/FRT/TO (V652020, Thermo Fisher Scientific). Although the human ribosomal protein L10a is referred to as uL1 in standard nomenclature (36), we used L10a here following the development of the TRAP method.

A homemade anti-ribosomal protein P0 antibody (9D5) was kindly provided by Dr. Toshio Uchiumi and Dr. Hiroe Sato (Niigata University) and is available from MBL (RN004M, MBL). The isotype control antibody for 9D5 (mouse IgG2a κ) was purchased from Proteintech (65208-1-IG). Antibodies against ribosomal proteins uL3 (GTX114725, GeneTex), L10a (A305-061A, BETHYL), uS2 (GTX114734, GeneTex), uS3 (GTX103964, GeneTex), uS15 (GTX101839, GeneTex), and PABP4 (A301-467A, BETHYL), CBP80 (A301-793A, BETHYL) and GAPDH (M171-7, MBL) were purchased commercially. A homemade anti-eIF4A3 polyclonal antibody was prepared as described previously (37).

### Cell lines and animals

Flp-In T-REx 293 cells (R78007, Thermo Fisher Scientific; mentioned as HEK293 in the text and figures) were cultured at 37ºC in Dulbecco’s modified Eagle’s medium (DMEM) supplemented with 10% fetal bovine serum, 100 U/mL penicillin, and 100 μg/mL streptomycin. pcDNA5/FRT/TO_EGFP-L10a was co-transfected with pOG44 (V600520, Thermo Fisher Scientific) into Flp-In T-REx 293 cells to generate a stable cell line inducibly expressing EGFP-L10a and selected in media supplemented with 50 μg/mL hygromycin according to the manufacturer’s instructions. EGFP-L10a expression was induced for 24 hours by the addition of doxycycline at 3 μg/mL. Zebrafish were grown at 28°C in E3 embryo medium (5 mM NaCl, 0.17 mM KCl, 0.33 mM CaCl_2_, and 0.33 mM MgSO_4_) using standard methods (38). *Drosophila* S2 cells were maintained in Schneider’s *Drosophila* medium (21720024, Thermo Fisher Scientific), supplemented with 10% fetal bovine serum and 1x antibiotic-antimycotic (15240096, Thermo Fisher Scientific) at 27ºC. Preparation of synchronized KH1668 *smg-2* (*yb979*) worms were prepared as previously described (39). Briefly, synchronized gravid worms were cultivated in S-complete medium supplemented with *E. coli* strain OP50 and bleached with a standard bleach solution (40). The embryos were harvested and washed three times with M9 buffer. The embryos were then incubated in M9 buffer for 18 h at 20ºC with gentle agitation for hatching, and L1 larvae were harvested and washed three times with M9 buffer.

### Immunoprecipitation of ribosome

HEK293 cells were seeded in a 6-well plate at 2.0 × 10^5^ cells per well. After 24 hours, the cells were lysed in 200 μL of lysis buffer (20 mM HEPES-NaOH pH 7.5, 2.5 mM MgCl_2_, 150 mM NaCl, 1% [v/v] Triton X-100, 0.5 mM DTT, protease inhibitor cocktail, phosphatase inhibitor cocktail, 100 μg/mL cycloheximide). The cell lysates were clarified by centrifugation at 20,000 x *g* at 4ºC for 10 min. The supernatant was mixed with 0.1 μg/μL RNase A or 1 unit/μL RNasin Plus and incubated with 1 μg of anti-ribosomal protein P0 antibody (isotype: mouse IgG2a κ, clone:9D5) (RN004M, MBL) or isotype control antibody (65208-1-IG, ProteinTech) at 4ºC for 60 min. 7 μL of Dynabeads Protein G (Invitrogen) was added to the mixture and incubated at 4ºC for 40 min. The beads were washed three times with 1 mL of wash buffer (20 mM HEPES-NaOH pH 7.5, 2.5 mM MgCl_2_, 150 mM NaCl, 0.05% [v/v] Tween 20) and directly mixed with 30 μL of SDS-PAGE sample buffer, heated at 95ºC for 5 min, and analyzed by western blotting. All proteins were detected using an ECL select western blotting detection reagent (RPN2235, cytiva) or ImmunoStar (296-69901, Wako) and imaged using a ImageQuant LAS 4000 mini (cytiva).

For immunoprecipitation of ribosomes from *Drosophila* S2 cells, 1.5 × 10^6^ cells were harvested by centrifugation at 600 x *g* for 5 min. The cell pellets were lysed in 400 μL lysis buffer (50 mM HEPES-NaOH pH 7.5, 10 mM MgCl_2_, 150 mM NaCl, 1% [v/v] NP-40, 0.5 mM DTT, protease inhibitor cocktail, phosphatase inhibitor cocktail, 100 mg/mL cycloheximide). The lysates were clarified by centrifugation at 20,000 x *g* for 10 min at 4ºC. The supernatant was divided into two fresh microcentrifuge tubes and incubated with 1 μg of 9D5 antibody or isotype control IgG at 4ºC for 60 min. The subsequent procedures were performed as described above.

For immunoprecipitation of the ribosome from zebrafish, 30 whole zebrafish embryos (4 days after fertilization) were resuspended in 1 mL lysis buffer (50 mM HEPES-NaOH pH 7.5, 100 mM KCl, 12 mM MgCl2, 1% [v/v] NP-40, 0.5 mM DTT, protease inhibitor cocktail, phosphatase inhibitor cocktail, 100 mg/mL cycloheximide, 1 mg/mL heparin, 0.2 unit/μL RNasin) and homogenized by douncing on ice. The lysates were then clarified by centrifugation at 10,000 x *g* at 4ºC for 10 min. The supernatant was divided into two fresh microcentrifuge tubes and incubated with 1 μg of 9D5 antibody or isotype control IgG at 4ºC for 60 min. Subsequent procedures were performed as described.

For immunoprecipitation of the ribosomes from the worms, synchronized L1 larvae were harvested by centrifugation at 600 x *g* for 1 min. The worm pellets were washed twice with 5 mL of M9 buffer and resuspended in 1 mL worm lysis buffer (50 mM HEPES-NaOH pH 7.5, 10 mM MgCl_2_, 150 mM NaCl, 1% [v/v] NP-40, 0.5 mM DTT, protease inhibitor cocktail, phosphatase inhibitor cocktail, 100 mg/mL cycloheximide). After homogenization by sonication, the lysates were clarified by centrifugation at 20,000 x *g* at 4ºC for 10 min. The supernatant was divided into two fresh microcentrifuge tubes and incubated with 1 μg of 9D5 antibody or isotype control IgG at 4ºC for 60 min. The subsequent procedures were performed as described above.

### ER stress and 4sU metabolic RNA labeling

The HEK293 cell line used for the inducible expression of EGFP-L10a was seeded in a 10 cm dish at 5.0 × 10^5^ cells per dish. The expression of EGFP-L10a was induced by 3 μg/mL of doxycycline for 24 h. On the next day, the medium was quickly replaced with a freshly prepared medium containing 200 μM 4sU with either 0.02% (v/v) DMSO or 1 μM thapsigargin. The cells were incubated at 37ºC for 3 hours and lysed in 1 mL lysis buffer (20 mM HEPES-NaOH pH 7.5, 2.5 mM MgCl_2_, 150 mM NaCl, 1% [v/v] Triton X-100, 0.5 mM DTT, protease inhibitor cocktail, phosphatase inhibitor cocktail, 100 mg/mL cycloheximide). Whole cell lysates were clarified by centrifugation at 10,000 x *g* at 4ºC for 10 min and the supernatant supplemented with 1 unit/μL RNasin Plus was used for further procedure. For cytosolic RNA preparation, 200 μL of the supernatant was mixed with 500 μL of ISOGEN II (311-07361, NIPPON GENE) supplemented with 1 mM DTT and the prepared sample solution was stored at -20ºC. For P-TRAP, 400 μL of the supernatant were incubated with 4 μg of 9D5 antibody at 4ºC for 60 min. Aftewards, 28 μL of Dynabeads Protein G was added to the mixture and incubated at 4ºC for 40 min. After the beads were washed three times with wash buffer (20 mM HEPES-NaOH pH 7.5, 2.5 mM MgCl_2_, 150 mM NaCl, 0.05% [v/v] Tween 20), the beads were directly mixed with 500 μL of ISOGEN II supplemented with 1 mM DTT and the sample solution was stored at -20ºC. For EGFP-L10a TRAP, 400 μL of the supernatant were incubated with 25 μL of GFP-Trap^®^ Magnetic Particles M-270 (gtd-100, proteintech) at 4ºC for 60 min. After the beads were washed three times with wash buffer, the beads were directly mixed with 500 μL of ISOGEN II supplemented with 1 mM DTT and the sample solution was stored at -20ºC. The manufacturer’s instructions were followed for RNA extraction using ISOGEN II, and the RNA samples were assessed for quality and quantity on MultiNA (Shimazu), a capillary electrophoresis instrument.

### SLAMseq and 3’ polyadenylated RNA-seq

Cytosolic and immunoprecipitated RNA were processed according to the standard SLAMseq protocol described previously (9). In brief, 5 μg of cytosolic RNA or immunoprecipitated RNA were incubated in 50 μL of alkylation buffer [50 mM NaPO4 buffer pH 8.0, 50% (v/v) DMSO, 10 mM iodoacetamide] at 50ºC for 15 min. The reaction was quenched by adding 1 μL of 1 M DTT, followed by ethanol precipitation. For each sample, 500 ng of cytosolic RNA or immunoprecipitated RNA was used as an input for QuantSeq 3’ mRNA-Seq Library Prep Kit FWD for Illumina (Lexogen). The cDNA library was prepared according to the manufacturer’s instructions. Libraries were assessed for quality using a MultiNA capillary electrophoresis instrument (Shimadzu), multiplexed to equimolar concentrations, and sequenced using the HiSeq 2500 system (Illumina).

### Bioinformatic analysis

Nascent and pre-existing polyadenylated RNAs were quantified based on sequence data using SlamDunk v 0.4.2, a pipeline for SLAMseq data analysis, with default parameters (9). Because QuantSeq targets mainly the 3’ end of individual RNAs in a poly(A) tail-dependent manner, the sequence data were aligned on genome-wide 3’ UTR sequences generated based on the human genome sequence (GRCh38.p13) and annotation data (gencode.v39.annotation.gff3). Twelve bases from the 5’ end were trimmed as adaptor-clipped reads, and then four or more subsequent adenines from the 3’ end were regarded as the remaining poly(A) tail and removed. Up to 100 regions with multiple mapped reads were allowed. Differential gene expression analysis was performed using DESeq2 v 1.3.8 (41). Differentially transcribed genes and differential translation efficiency genes (FDR < 0.05) were identified using the deltaTE method (42), with read counts generated by SlamDunk using cytosolic RNA-seq and P-TRAP-seq as inputs. Gene ontology analysis was performed by over-representation analysis using a biological process database (WebGestalt 2019) (43).

## Results

### Combination of TRAP with 4sU metabolic RNA labeling for simultaneous measurement of nascent transcripts and their translation

SLAMseq distinguishes 4sU-labeled nascent transcripts from pre-existing RNA prior to labeling by bioinformatics detection of specific T>C conversions at the sites of 4sU incorporation. The combination of SLAMseq with QuantSeq, which analyzes the inherently uridine-rich 3’ UTRs of polyadenylated RNAs, allows the robust quantification of T>C conversions (9). As QuantSeq requires reverse transcription from poly(A), it is compatible with polysome profiling and TRAP techniques for recovering polyadenylated RNAs bound to ribosomes. Polysome profiling allows for the fractionation of monosomes and polysomes using a sucrose density gradient to monitor active translation. However, it requires specialized equipment (e.g., ultracentrifuge and gradient fractionation systems), is labor-intensive, and does not allow the handling of many samples in parallel (24). To ensure experimental throughput and to measure nascent transcripts and translation, we designed a combination of TRAP and SLAMseq/QuantSeq (Figure 1).

**Figure 1.**
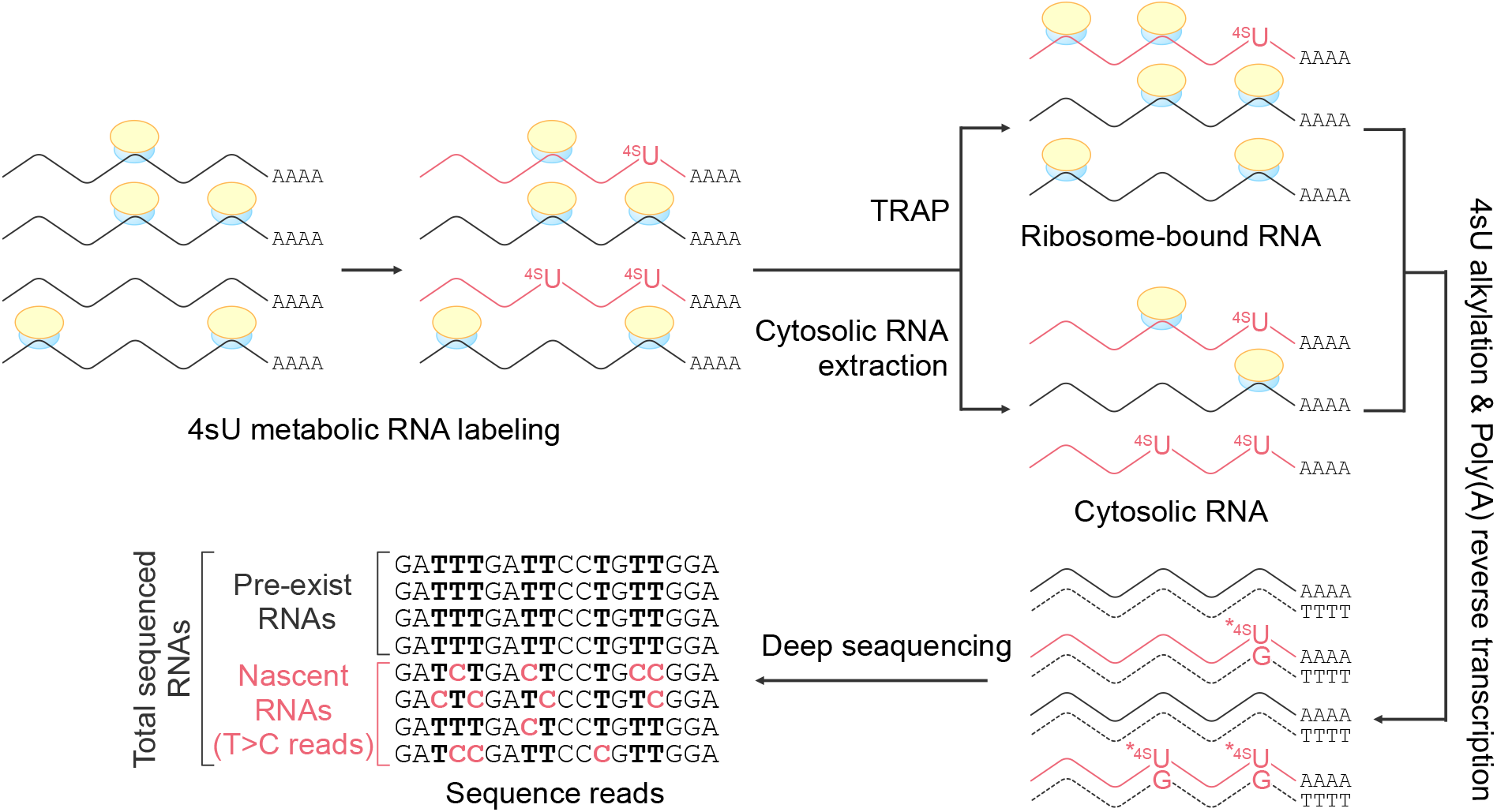
Simultaneous measurement of transcriptome and translatome by combining TRAP with 4sU metabolic RNA labeling. Schematic representation of the procedure for combing 4sU metabolic RNA labeling using the TRAP technique. Cultured cells are incubated with 4-thiouridine (4sU) for a defined period of time. Cells are lysed in a lysis buffer containing cycloheximide to prevent ribosome run-off and divided into cytosolic RNA and TRAP fractions. RNAs are reverse transcribed from poly(A) tail according to QuantSeq (3’ mRNA sequence) for cDNA library preparation and deep sequencing. Nascent transcripts are distinguished from whole RNAs by the detection of T>C conversion at sites of 4sU incorporation.

### Affinity purification of endogenous translating ribosome from human cultured cells using antibody against ribosomal P-stalk

TRAP requires genetically engineered cells expressing affinity-tagged ribosomal proteins of the large ribosomal subunit to purify translating ribosome-bound RNAs. The expression of affinity-tagged ribosomal proteins with specific promoters enables the measurement of cell-, tissue-, and developmental stage-specific translatome (22,27-29) which is a major advantage of TRAP. However, the genetic manipulation of endogenous ribosomal proteins reduces experimental throughput and is unsuitable for cells that cannot be genetically engineered. Therefore, we aimed to develop a high-throughput system that allows affinity purification of endogenous non-tagged ribosomes from a variety of cells and organisms. To this end, we focused on the ribosomal P-stalk, a multimeric ribosomal protein complex in the eukaryotic 60S ribosomal subunit. The P-stalk consists of one copy of ribosomal protein P0 and two heterodimers of ribosomal proteins P1 and P2, forming a pentametric P0-(P1-P2)_2_ complex (44). P0, P1, and P2 share homologous C-terminal intrinsically disordered regions (IDRs); therefore, the P-stalk contains five copies of C-terminal IDRs (45). Because the homologous C-terminal IDRs of the P-stalk are exposed outside the ribosome, the antibody that recognizes the C-terminal IDRs of the P-stalk is expected to be suitable for the immunoprecipitation of endogenous ribosomes (Figure 2A) (46). Metz et al. recently applied an endogenous ribosome immunoprecipitation approach using 5.8S rRNA as an epitope (47). Our method uses multiple epitopes in the ribosome instead of a single epitope for efficient immunoprecipitation.

**Figure 2.**
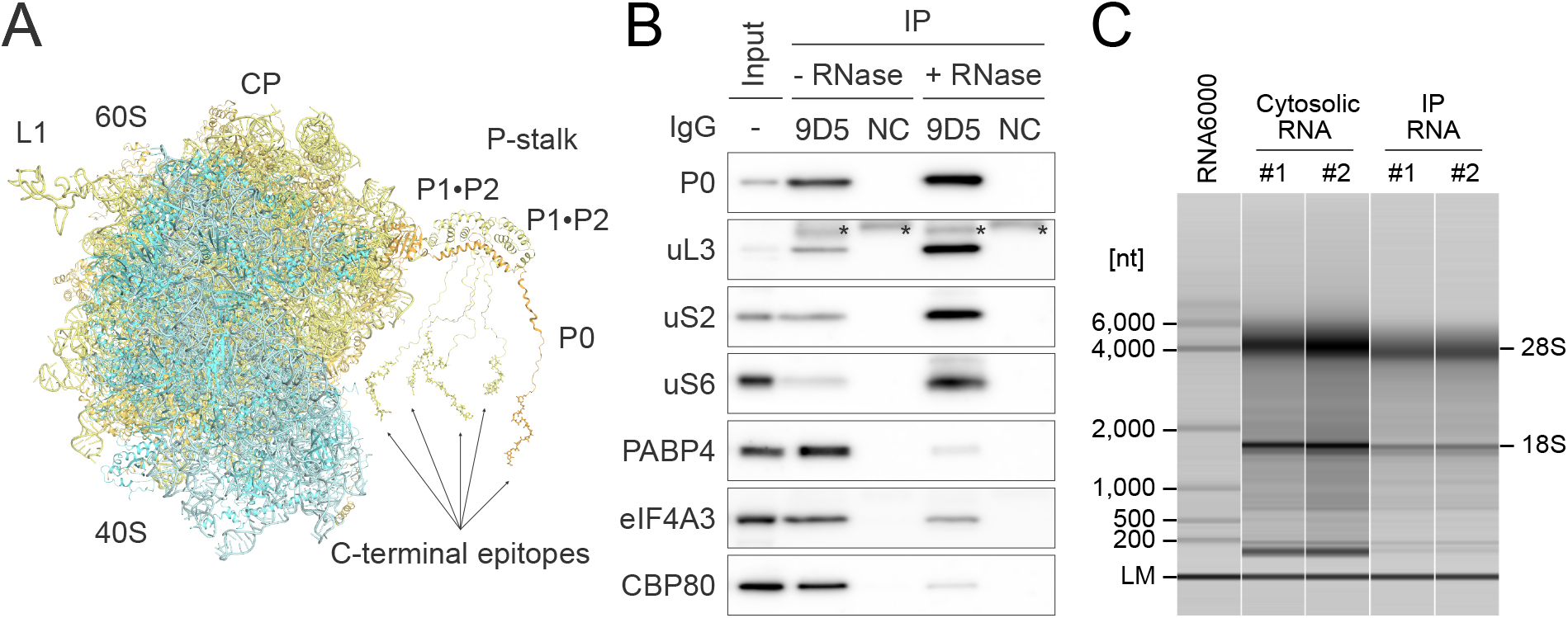
Immunoprecipitation of endogenous translating ribosome mediated by the C-terminal IDRs of the P-stalk. (A) A structural model of human 80S ribosome containing ribosomal P-stalk generated from three coordinates [Protein data bank code: 3A1Y, 4BEH, and 6EK0 (36,52,53)]. The C-terminal IDR of P0 is modeled arbitrarily. The epitopes of the anti-P0 antibody (9D5) are shown as a stick model. Colors are as follows: 40S ribosomal subunit (cyan), 60S ribosomal subunit (pale yellow), and ribosomal protein P0 (orange). (B) Immunoprecipitation of the endogenous ribosome and ribosome-bound RNA with the anti-ribosomal protein P0 antibody (9D5) or isotype control IgG (NC) from the cytosolic lysate of HEK293 cell in the presence or absence of RNase A, followed by western blotting. The input contained 1% of the lysate used for immunoprecipitation. Asterisks (*) indicate non-specific signals from the antibodies (9D5 or NC). (C) Examples of capillary electrophoresis profiles for cytosolic RNAs and immunoprecipitated RNAs from HEK293 cells. Two independent experiments were performed for each condition (#1, #2). The lower marker (LM) indicates internal standards (25 nt). RNA ladder (the Agilent RNA 6000 Pico kit) was used as a size marker.

We tested whether translating ribosomes could be immunoprecipitated from cultured human cell extracts, using a monoclonal antibody (9D5) that recognizes the C-terminal IDRs of P0, P1, and P2 (48). The isotype control of 9D5 was used as a negative control for immunoprecipitation. As expected, the 9D5 antibody immunoprecipitated with ribosomal proteins uL3, uS2, and uS6, and mRNA-binding proteins CBP80, eIF4A3, and PABP4 (Figure 2B). RNase A treatment markedly decreased the intensity of CBP80, eIF4A3, and PABP4 but not that of ribosomal proteins, revealing that these mRNA-binding proteins were co-immunoprecipitated via ribosome-bound mRNA. In addition, the RNA electrophoresis profiles confirmed that representative ribosomal RNA signals (28S and 18S) and smear patterns of ribosome-bound RNAs were obtained by co-immunoprecipitation (Figure 2C). These results indicate that the 9D5 antibody against the ribosomal P-stalk enables the purification of endogenous non-tagged ribosomes and ribosome-bound RNAs.

### Affinity purification of endogenous ribosome via ribosomal P-stalk from diverse eukaryotes

A C-terminal 22 amino acids sequence in the IDRs of the P-stalk recognized by the 9D5 antibody is well-conserved in diverse eukaryotes from yeast to humans (Figure 3A). Therefore, the immunoprecipitation of endogenous ribosomes using the 9D5 antibody may be applied to diverse eukaryotes. We next tested whether the immunoprecipitation of endogenous ribosomes using the 9D5 antibody could be performed in zebrafish, fruit flies, and worms (Figures 3B–D). In these experiments, the 9D5 antibody immunoprecipitated with the ribosomal proteins uL3, uS3, and uS15, indicating that the 9D5 antibody can be used to purify endogenous ribosomes from diverse eukaryotes without genetic manipulation. We termed the anti-P0 antibody-mediated purification of endogenous ribosomes P-stalk-mediated translational ribosome affinity purification (P-TRAP).

**Figure 3.**
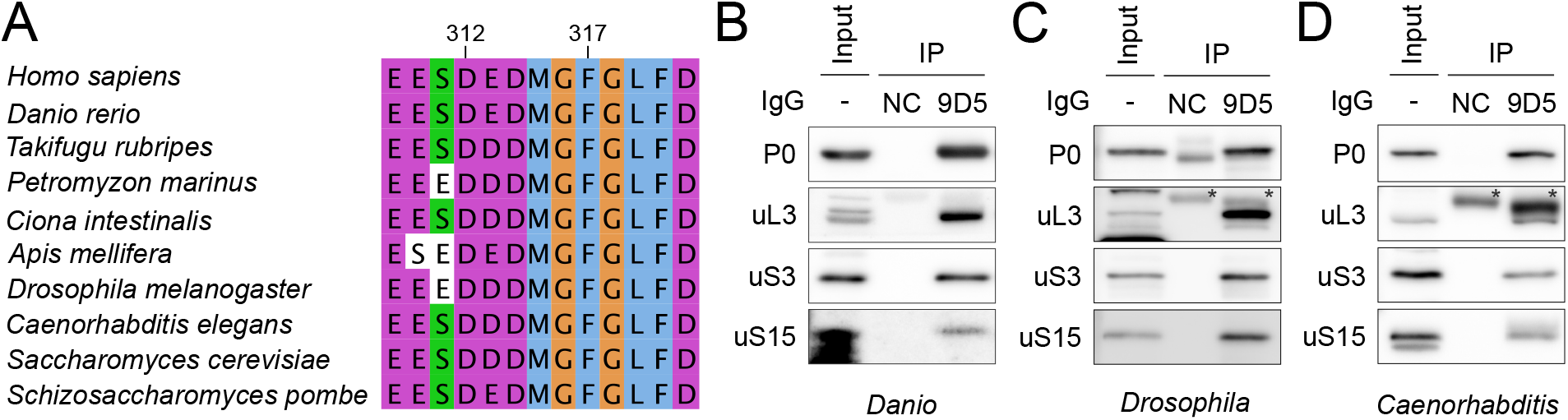
Immunoprecipitation of endogenous ribosome from diverse eukaryotes mediated by the C-terminal IDRs of the P-stalk. (A) Sequence alignment of the C-terminal 12 amino acids of ribosomal protein P0 in diverse eukaryotes. The 9D5 anti-P0 antibody recognizes these sequences. (B–D) Immunoprecipitation of the endogenous ribosome with the 9D5 antibody or isotype control IgG (NC) from whole zebrafish embryo (B), *Drosophila* S2 cells (C), and adult *C. elegans* (D) in the presence of RNase A, followed by western blotting. The input contained 1% of the lysate used for immunoprecipitation. Asterisks (*) indicate non-specific signals from the antibodies (9D5 or NC).

### Simultaneous measurement of nascent transcriptome and translatome in the UPR by combining SLAMseq and QuantSeq with P-TRAP or conventional TRAP

We used a combination of SLAMseq/QuantSeq with P-TRAP to analyze nascent polyadenylated RNAs and their translation at the level of individual genes in the unfolded protein response (UPR), whereby transcription and translation are dynamically reprogrammed to reduce the loading of unfolded or misfolded proteins into the endoplasmic reticulum (ER) and restore protein homeostasis (49,50). To compare P-TRAP with conventional TRAP, we generated a HEK293 cell line for the stable expression of doxycycline-inducible enhanced green fluorescent protein (EGFP) fused with the N-terminus of ribosomal protein L10a (EGFP-L10a), an initially reported TRAP technique in mammals (Figure S1A) (22,27-29). The expression of EGFP-L10a enabled the affinity purification of translating ribosomes containing EGFP-L10a using GFP nanobody-conjugated magnetic beads (Figure S1B, S1C). We subjected the EGFP-L10a-expressing cells with 4sU for metabolic RNA labeling and thapsigargin, which inhibits the sarcoplasmic/endoplasmic reticulum calcium ATPase (SERCA) and leads to the accumulation of unfolded proteins in the ER for 3 hours (51). The cell lysate was divided into three factions: cytosolic RNA (fraction I), ribosome-bound RNAs obtained by P-TRAP (fraction II), and conventional EGFP-L10a-TRAP (fraction III), followed by the 3’ polyadenylated RNA-seq (here we termed cytosolic RNA-seq, P-TRAP-seq, and L10a-TRAP-seq). Principal component analysis (PCA) revealed that cytosolic RNA-seq, P-TRAP-seq, and L10a-TRAP-seq were distinguishable in the DMSO- and thapsigargin-treated groups (Figure S2). In particular, the cytosolic RNA group was separated from the other two groups in PC1 (41.0% and 38.9% for total and T>C reads, respectively), reflecting a low correlation between the transcriptome characterized by standard RNA-seq and the translatome (4,19-21) both in the presence or absence of thapsigargin.

### Translatome analysis in the UPR by P-TRAP-seq and conventional TRAP-seq

To validate whether translatome analysis with P-TRAP recapitulated gene expression reprogramming in the UPR, we performed differential gene expression analysis using total reads (Figure 4A) to identify differentially expressed genes (DEGs). Compared to cytosolic RNA-seq, more DEGs were detected in P-TRAP-seq and L10a-TRAP-seq (614 genes in cytosolic RNA-seq, 894 genes in P-TRAP-seq, and 807 genes in L10a-TRAP-seq). Similar to cytosolic RNA-seq and conventional L10a-TRAP-seq, the expression of typical genes involved in the UPR (*HSPA5, DDIT3, PDIA4, DNAJC2*, and *HERPUD1*) was significantly increased in P-TRAP-seq analysis (Figure 4B). Notably, in contrast to genes regulated by RNA, specific genes preferentially translated by polysomes in the UPR obtained by sucrose density gradient and DNA microarray (52) were upregulated in the P-TRAP-seq and L10a-TRAP-seq data, but not in the cytosolic RNA-seq data (Figure 4C). These results indicate that P-TRAP-seq detects changes in translation without changing the RNA content and is well suited for translatome analysis like conventional TRAP method and polysome profiling.

**Figure 4.**
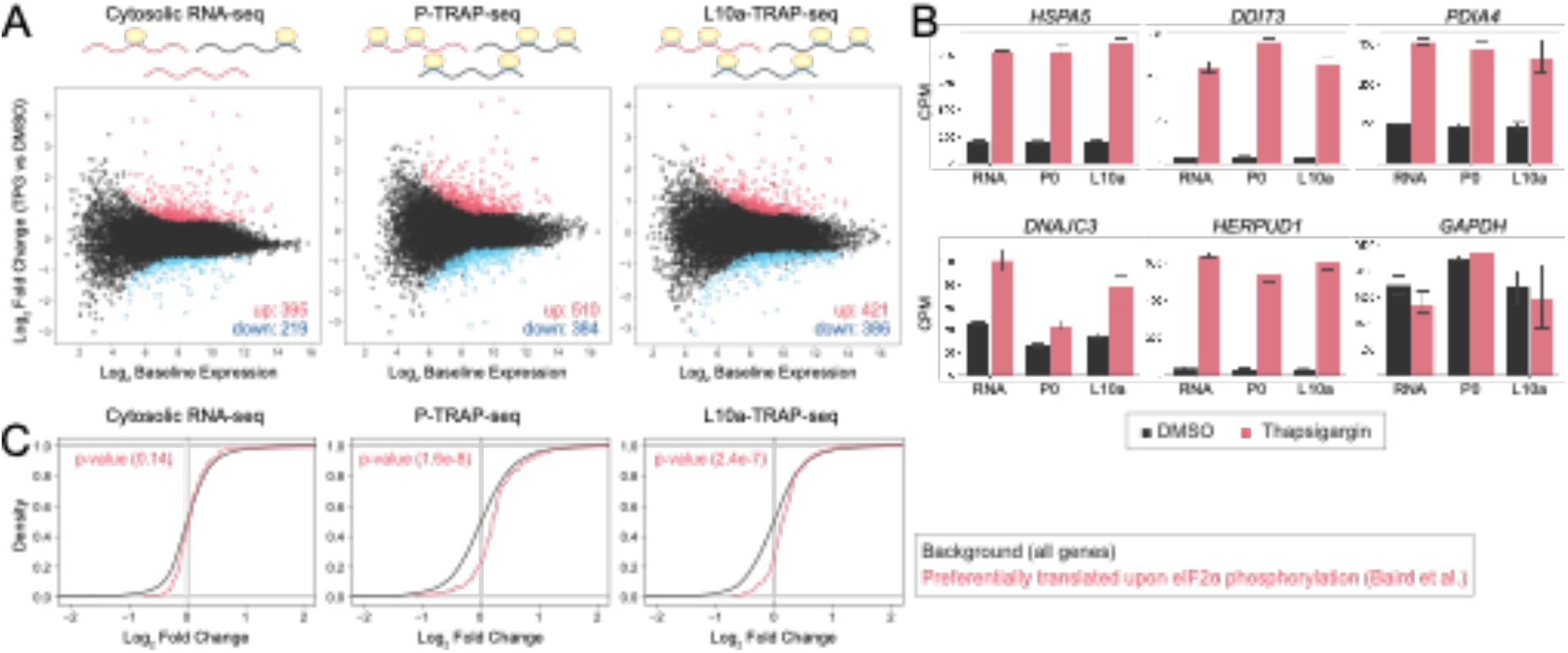
Transcriptome and translatome changes in the UPR at the levels of whole transcripts analysis. (A) MA plot of cytosolic RNA-seq, P-TRAP-seq, and L10a-TRAP-seq data from TPG-treated versus DMSO-treated HEK293 cells. Differentially expressed genes (adjusted p-value < 0.01 and log2 fold change > 1.5) are highlighted in red (up-regulated) or blue (down-regulated). (B) CPM (counts per million) values of selected ER stress response genes (*HSPA5, DDIT3, PDIA4, DNAJC3*, and *HERPUD1*) and a control gene (*GAPDH*) in the cytosolic RNA-seq (RNA), P-TRAP-seq (P0), and L10a-TRAP-seq (L10a) in response to DMSO-(black) or TPG-(red) treatments. The averages and standard deviations of three replicates are shown. (C) Cumulative distributions of the log2 fold changes of selected gene categories (preferentially translated by polysome upon eIF2α phosphorylation) compared to all genes, together with p-values of Mann-Whitney U test.

### Categorical analysis of nascent transcriptome and translatome in the UPR

In SLAMseq, 4sU incorporated into nascent RNA is alkylated during library preparation, and T>C conversion occurs via reverse transcription. Consistent with a previous report (9), we observed a median rate of 2.2–2.8% for T>C conversion and <0.2% for any other conversion in total transcripts in the presence of 4sU (Figure S3). We extracted sequence reads with T>C conversion using the SlamDunk pipeline and obtained the nascent transcriptome (cytosolic RNA-seq) and translatome (P-TRAP-seq and L10a-TRAP-seq) during the UPR. Differential gene expression analysis of nascent RNAs revealed 685, 694, and 540 DEGs in the cytosolic RNA-seq, P-TRAP-seq, and L10a-TRAP-seq, respectively (Figure 5A). Similar to the total reads analysis, the expression of typical genes involved in the UPR increased in the nascent transcriptome and translatome (Figure 5B). SLAMseq has been shown to increase the detection power of DEGs by focusing on actively transcribed RNAs (16). Consistent with this, SLAMseq of cytosolic RNA during the acute UPR increased the number of DEGs by focusing on nascent RNAs (685 DEGs in T>C reads and 614 DEGs in total reads). Notably, the DEGs detected in the translatome analysis (P-TRAP-seq and L10a-TRAP-seq) were higher for whole RNAs than for nascent RNAs (694 DEGs in T>C reads and 894 DEGs in total reads in P-TRAP-seq, 540 DEGs in T>C reads, and 807 DEGs in total reads in L10a-TRAP-seq). These trends suggest that a major part of the translatome changes in the early UPR are due to the alteration in translation efficiency of pre-existing RNAs.

**Figure 5.**
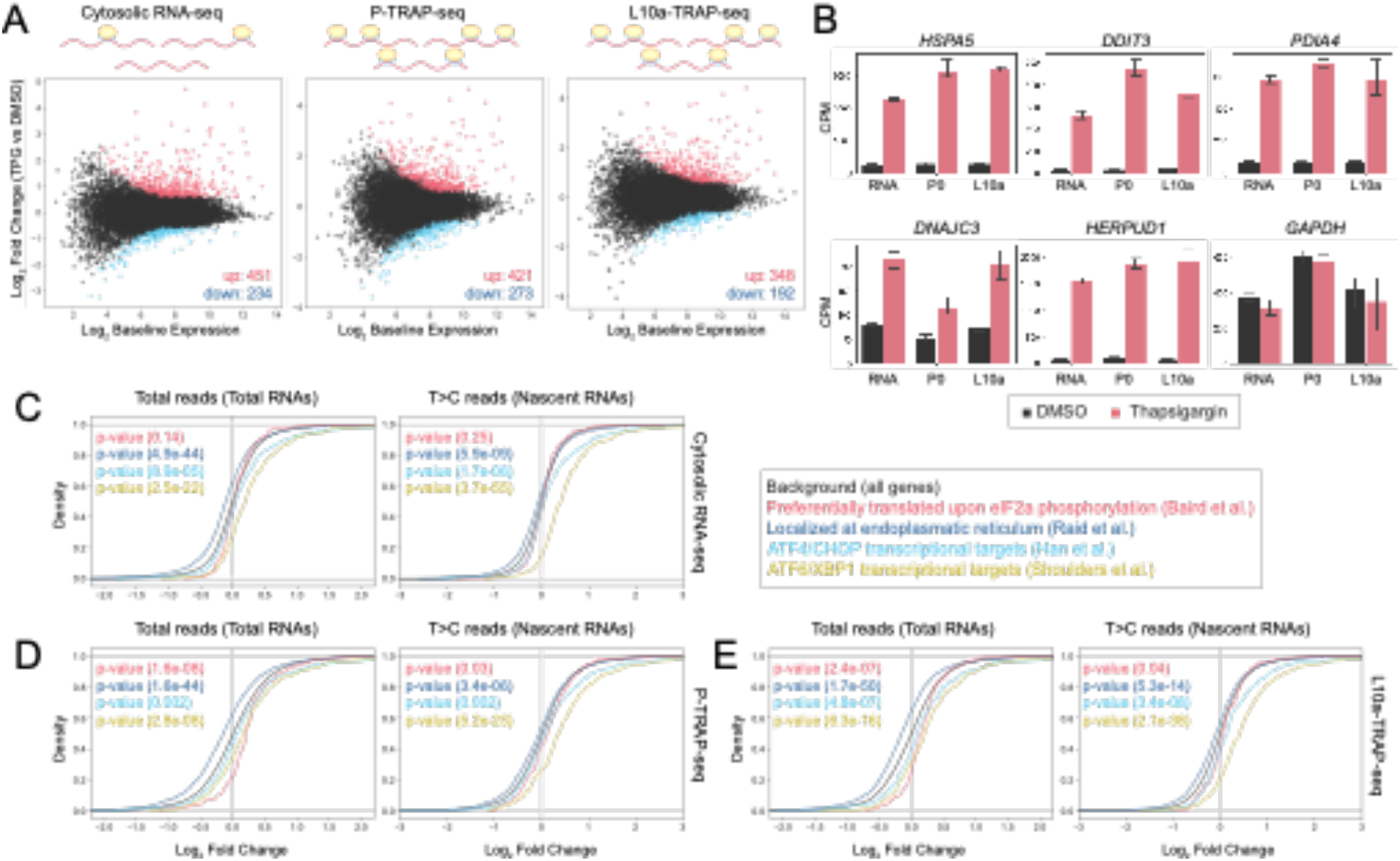
Transcriptome and translatome changes in the UPR at the levels of nascent transcripts analysis. (A) MA plot of cytosolic RNA-seq, P-TRAP-seq, and L10a-TRAP-seq data from TPG-treated versus DMSO-treated HEK293 cells. Differentially expressed genes (adjusted p-value < 0.01 and log_2_ fold change > 1.5) are highlighted in red (up-regulated) or blue (down-regulated). (B) CPM (counts per million) values of selected ER stress response genes (*HSPA5, DDIT3, PDIA4, DNAJC3*, and *HERPUD1*) and a control gene (*GAPDH*) in the cytosolic RNA-seq (RNA), P-TRAP-seq (P0), and L10a-TRAP-seq (L10a) in response to DMSO-(black) or TPG-(red) treatments. The averages and standard deviations of three replicates are shown. (C) Cumulative distributions of the log_2_ fold changes of selected gene categories (preferentially translated by polysome upon eIF2α phosphorylation, localized at endoplasmic reticulum, ATF4/CHOP transcriptional targets, and ATF6/XBP1 transcriptional targets) compared to all genes, together with p-values of Mann-Whitney U test.

In the acute phase UPR, the ER stress sensor PERK phosphorylates translation initiation factor 2α (eIF2α), leading to global repression of translation initiation, and the whole protein synthesis rate is transiently reduced to 15–20% (53,54). The phosphorylation of eIF2α also promotes the preferential translation of mRNAs with upstream open reading frames (uORFs), such as transcription factors ATF4 and CHOP. As described in Figure 4C, the RNAs enriched in the polysome fraction upon eIF2α phosphorylation increased translation levels in both P-TRAP-seq and L10a-TRAP-seq in whole RNAs analysis. In contrast, the translation levels of these RNAs were not significantly elevated in the nascent translatome suggesting that the phosphorylated eIF2α dependent preferential translation mainly occurs to pre-existing, but not to nascent RNAs in the UPR (Figure 5C–E: red lines). We then analyzed gene expression reprogramming during the UPR for several gene categories (Figure 5C), such as ATF4 and CHOP (ATF4/CHOP targets: cyan lines) and another essential branch in the UPR, which includes ATF6 and XBP1 (ATF6/XBP1 targets: yellow lines). Consistent with previous reports (55,56), the RNA levels of ATF4/CHOP and ATF6/XBP1 targets were significantly elevated upon thapsigargin treatment, which was particularly prominent for the target genes of ATF6/XBP1 in the nascent transcriptome (Figure 5C). These results are consistent with the enhanced transcriptional activity of ATF4/CHOP and ATF6/XBP1 during the UPR. Abundance of ATF6/XBP1 targets was also significantly elevated in nascent translatomes obtained by P-TRAP-seq and L10a-TRAP-seq, indicating that the nascent RNAs transcribed by ATF4/CHOP and ATF6/XBP1 were efficiently translated during the UPR (Figure 5D and 5E).

In the acute phase of the UPR, the total amount of proteins synthesized on the endoplasmic reticulum membrane is reduced to decrease protein loading to the ER (49,50). The expression of genes encoding membrane and secretory proteins was significantly downregulated in both total reads and T>C reads in the cytosolic RNA-seq data in response to ER stress (Figure 5C–E: blue lines). Notably, the reduction in RNA levels localized to the ER membrane was more prominent in the total reads than in the T>C reads. Similar patterns were observed in the P-TRAP-seq and L10a-TRAP-seq data. These results indicate that the UPR likely destabilizes pre-existing RNAs localized to the ER membrane, known as the output of regulated IRE1-dependent decay (RIDD) (57,58).

### Analysis of nascent transcriptome and translatome in the UPR for individual genes

The advantage of combining TRAP-seq and SLAMseq/QuantSeq is the robust estimation of expression and translation levels of nascent polyadenylated RNAs for individual genes. To clarify the translational states in the UPR, we performed deltaTE analysis (42) for pre-existing or nascent RNAs from cytosolic RNA-seq and P-TRAP-seq data (Figure 6A). This analysis enabled us to identify (1) genes regulated at the RNA level (forward), (2) genes regulated at the translation level (exclusive), (3) genes regulated by the synergistic effect of RNA and translation levels (intensified), and (4) genes regulated by the levels of RNA and translation against each other (buffered). Consequently, deltaTE analysis revealed that the actively transcribed RNAs during UPR (categorized as forwarded) were loaded to the ribosomes indicating that these actively transcribed RNAs relatively escaped from the global repression of translation initiation by eIF2α phosphorylation. Actively translated RNAs (categorized as forwarded) included typical genes involved in the UPR (*HSPA5, HERPUD1*, and *PDIA4*), and enrichment analysis showed a significant enrichment of genes responsible for ER stress tolerance (Figure 6B). This data is consistent with previous observations that the UPR leads to massive translational repression associated with the phosphorylation of eIF2α, followed by synthesis of proteins involved in the proper folding or degradation of unfolded and misfolded proteins in the ER (59,60). When the UPR proceeds, phosphorylation of eIF2α is partially reduced, and global translation levels are gradually restored (53,54). Partial restoration of translational activity is important for protein synthesis from nascent RNAs in the UPR. Notably, deltaTE analysis revealed that a certain subset of nascent RNAs still undergo translational repression, although these RNAs are involved in the escape of ER stress tolerance (Figure 6A). The data suggests the existence of unknown mechanisms, distinct from eIF2α phosphorylation, to escape or to be translationally repressed. Consistent with this idea, a recent study revealed that the length of the poly(A) tail is an important determinant of the translation levels of XBP1s mRNA (61) in the chronic UPR. Further analysis is required to understand the complexity of reprogramming gene expression in the UPR.

**Figure 6.**
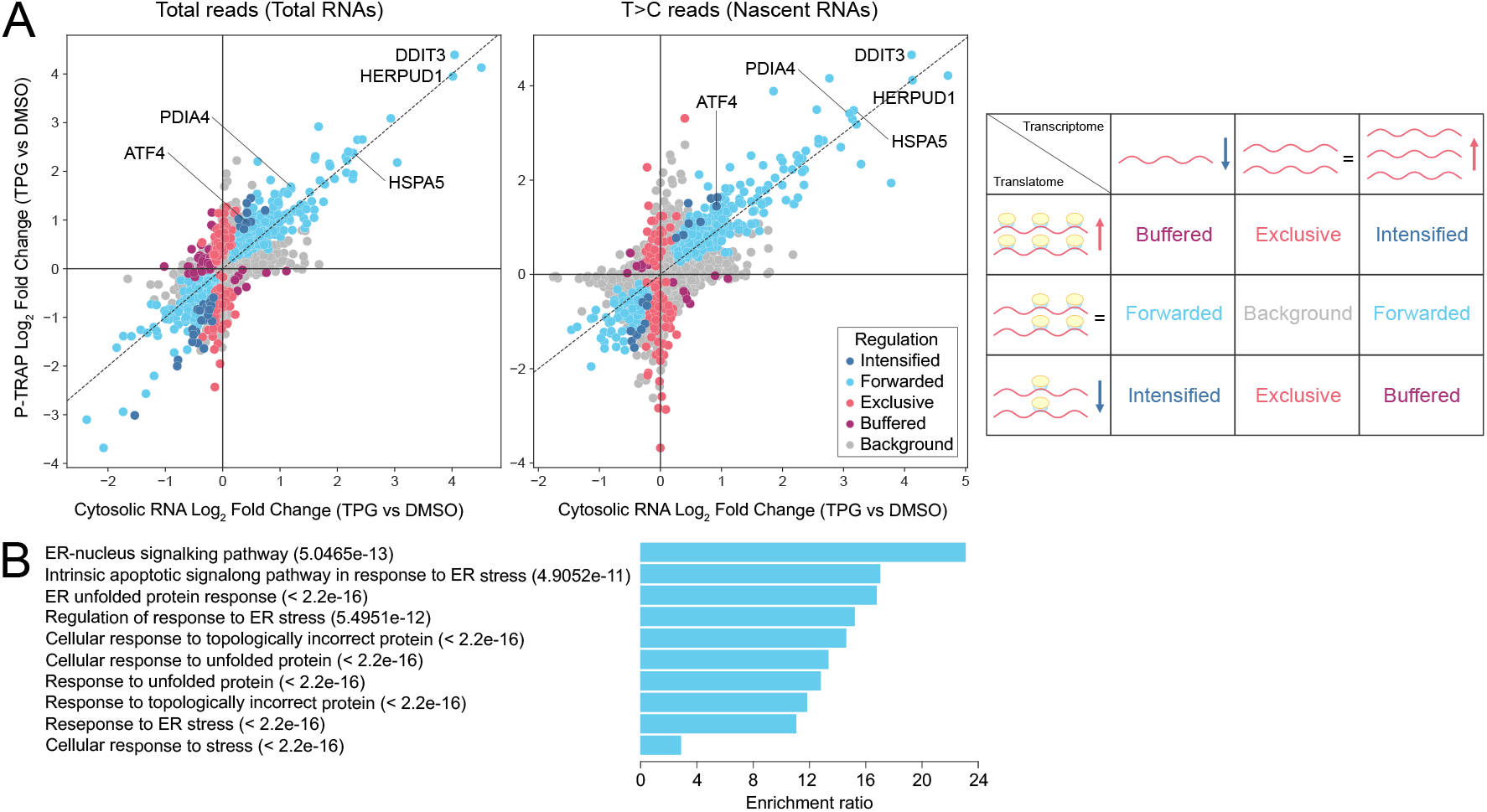
Translation regulation of nascent transcripts in the UPR at the level of individual genes. (A) Left: fold changes each total or nascent RNA in cytosolic RNA-seq and P-TRAP-seq data at the levels of the individual genes analyzed by the deltaTE method (42). Translationally forwarded genes (cyan), intensified genes (blue), exclusive genes (red), and buffered genes (purple) are highlighted. Right: Regulation modes categorized by deltaTE analysis. Translationally forwarded genes significantly changed in transcriptome and translatome at the same rate. Translationally exclusive genes have a significant change in translatome, with no change in the transcriptome. Translationally buffered have a significant change in translatome that counteracts the change in transcriptome; hence, buffering the effect of transcription. Translationally intensified genes have a significant change in translatome that acts with the effect of transcription. (B) The enrichment ratios in over-representation analysis (ORA) of up-regulated forward genes in deltaTE analysis (T>C reads) performed by WebGestalt 2019 analysis (43). Resulted biological processes and FDR values are shown.

## Discussion

In this study, we established a simple protocol for the simultaneous measurement of nascent polyadenylated RNAs and their translation at the gene level. Recently, Schott et al. developed nRibo-seq for the simultaneous measurement of nascent RNA synthesis and translation efficiency at the level of bulk RNAs or specific gene categories by combining Ribo-seq with 4sU metabolic RNA labeling (35). Instead of Ribo-seq, we combined TRAP-seq with 4sU metabolic RNA labeling. Because TRAP-seq is compatible with QuantSeq, enabling the accurate quantification of T>C conversions from a small number of sequence reads, it is suitable for the simultaneous measurement of nascent RNA synthesis and translation efficiency at the levels of individual genes. In addition, TRAP-seq does not require specialized equipment such as ultracentrifugation and gradient fractionation systems, is not labor-intensive, and enables the recovery of ribosome-bound RNAs from a small number of cells. These advantages of TRAP-seq make it compatible with automated library preparation systems for high-throughput sequencing procedures, such as single-cell RNA sequencing. Furthermore, in contrast to conventional TRAP-seq, which requires genetic manipulation for the affinity purification of ribosomes, the newly developed P-TRAP-seq method can be applied to immunoprecipitate endogenous ribosomes without genetic manipulation. Therefore, measuring nascent polyadenylated RNAs and their translation using P-TRAP-seq with 4sU metabolic RNA labeling is a simple and robust method for investigating the crosstalk between transcriptional and translational regulation. Although individual translation efficiency can be monitored using mass-spectrometry-based techniques such as PUNCH-P (62) and BONCAT (63), our sequencing-based method is simple, low-cost, and provides highly accurate measurements.

It should be noted that the translatome measured using TRAP-seq only reflects ribosome-bound RNA levels. In contrast to Ribo-seq, TRAP-seq cannot determine the position of individual ribosomes in RNAs and has less resolution for complex translational regulation (e.g., translation efficiency for multiple open reading frames in a single transcript). TRAP-seq relies on a simple immunoprecipitation technique to extract ribosomes, including monosomes and polysomes; therefore, it cannot selectively fractionate actively translating polysomes, similar to the polysome profiling technique (64). Since the polysome profile also yields full-length ribosome-bound RNA and is compatible with 3’ mRNA-seq, a combination of polysome profile and 4sU metabolic RNA labeling is preferred when analyzing more efficiently translated RNAs (i.e., RNAs bound to multiple ribosomes).

There are two major advantages to the immunoprecipitation of endogenous ribosomes with C-terminal IDRs of the ribosomal P-stalk. The first is the high specificity and efficiency of immunoprecipitation. The C-terminal IDRs of the P-stalk were present in five copies on the 60S large ribosomal subunit and exposed on the solvent side (Figure 2A). Although these IDRs bind to multiple translational GTPase factors (trGTPases), such as eEF1A and eEF2 (65), the 9D5 antibody binds preferentially and stably to the P-stalk rather than to trGTPases (48,66). In addition, because P-stalk incorporation into the pre-60S ribosomal subunit occurs in the late cytoplasmic stage of ribosome assembly, P-TRAP only captures mature translating ribosome particles (67,68). Therefore, antibodies recognizing the C-terminal IDRs of the P-stalk ensure stable, specific, and robust immunoprecipitation of endogenous translating ribosomes.

Secondly, the C-terminal IDRs of the P-stalk are conserved across most, if not all, eukaryotes (46). Indeed, antibodies that recognize the highly conserved C-terminal IDR allow the purification of endogenous ribosomes from cultured human cells, zebrafish, flies, and nematodes without any genetic manipulations (Figure 3). P-TRAP can shorten the experimental period compared with conventional TRAP, which requires genetic manipulation. Furthermore, immunoprecipitation of endogenous ribosomes in P-TRAP allows for translatome analysis from non-model organisms without the need for genetic manipulation techniques, or from human specimens in clinical research. Moreover, the immunoprecipitation of ribosomes using P-stalks can be applied to other experimental systems that require ribosome collection, such as Ribo-seq. Highly efficient ribosome collection without ultracentrifugation is expected to significantly reduce the time and effort required for experiments and simplify protocols. These advantages of P-TRAP will accelerate translatome research and contribute to our understanding of gene expression regulation in various biological fields such as UPR (49,50), nonsense-mediated mRNA decay (69), immune responses (70), and drug responses (44).

Recent proteomic analyses have shown diversity in the composition of ribosomal proteins, referred to as ribosomal heterogeneity, resulting in the preferential translation of specific RNA. For example, L10a is present at substoichiometric levels in mouse embryonic stem cells (mESCs), and ribosomes containing L10a have an altered affinity for some transcripts (71). Performing TRAP-seq using substoichiometric ribosomal proteins as bait for immunoprecipitation may cause translatome bias. The P-stalk is present in polysomes with 1:1 stoichiometry in mESCs, indicating that the translatome obtained with P-TRAP-seq would have less translation bias due to ribosome heterogeneity, making the P-stalk an ideal bait for the TRAP-seq technique. However, in several experimental designs, P-TRAP has a disadvantage compared to the conventional TRAP technique because of the purification of ribosomes without genetic manipulation. Conventional TRAP captures gene expression from particular cells/tissues of genetically modified organisms expressing affinity-tagged ribosomal proteins using cell/tissue-specific promoters, such as the Gal4-UAS system in the fly or the Cre-lox system in mice (27,28,72). In contrast to conventional TRAP, P-TRAP cannot fractionate ribosomes from specific cells or tissues. Therefore, it is essential to use conventional TRAP and/or P-TRAP according to the experimental design.

## Data availability

The deep sequencing data reported in this study have been deposited in the DDBJ (DRA015587). All codes necessary to perform the computational analysis are available at GitHub (https://github.com/hirosome/2023-trap-upr).

## Funding

This work was supported by JSPS KAKENHI (20J00036 and 21K15015 to H.I., 20J00036 and 20H03438 to A.Y., 20H04839 and 20H03181 to H.K.) and the Japan Science and Technology Agency (JPMJMS2022 to A.Y.). This work was also supported by research grants from the Manufacturing Promotion Division, Department of Commerce, Industry, and Labor, Okinawa Prefectural Government, Takeda Science Foundation, and Bristol Myers Squibb K.K..

## Conflicts of interest

The authors declare that they have no competing interests.

## Acknowledgments

We thank Dr. Toshio Uchiumi and Dr. Hiroe Sato for providing anti-ribosomal protein P0 antibodies. We thank Dr. Eriko Matsuura, Dr. Shintaro Iwasaki, and Dr. Yukihide Tomari for providing *Drosophila* S2 cells. We thank Ms. Naima Batool for editing a draft of the manuscript.

## Notes

### Competing Interest Statement

The authors have declared no competing interest.

