## Supplemental Information for "Simultaneous measurement of nascent transcriptome and translatome using 4-thiouridine metabolic RNA labeling and translating ribosome affinity purification"

*Corresponding authors:

### Supplementary Tables

**Table. S1.** Results of deferential gene expression analysis by DESeq2 for RNA-seq, P-TRAP-seq, and L10a-TRAP-seq.

**Table. S2.** Results of differentially transcribed genes and differentially translated genes analysis using deltaTE method for RNA-seq, P-TRAP-seq, and L10a-TRAP-seq.

### Supplementary Figures

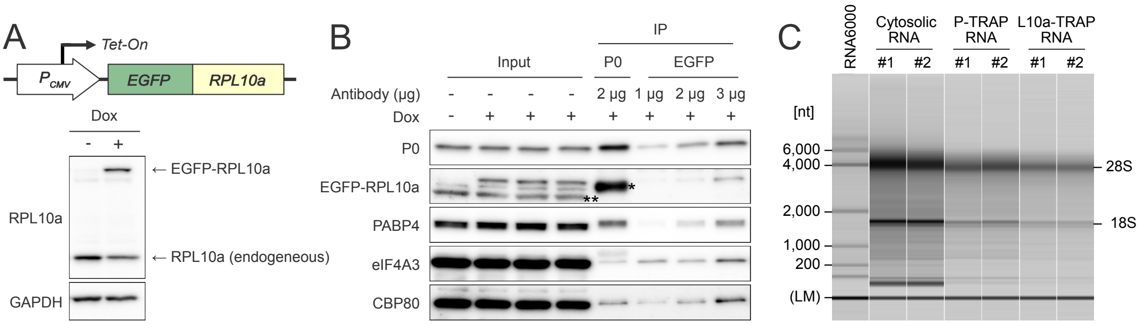


**Fig. S1.** (A) Top: schematic representation of HEK293 cell line stably expressing doxycycline-inducible (EGFP-RPL10a). Bottom: the doxycycline-inducible protein expression level of EGFP-RPL10a analyzed by western blotting. EGFP-RPL10a were detected by anti RPL10a antibody. (B) Immunoprecipitation of the endogenous ribosome and RNA binding proteins with the anti-ribosomal protein P0 antibody (9D5) or GFP-Trap® Magnetic Particles from the cytosolic lysate of HEK293 cell, followed by western blotting. The input contained 1% of the lysate used for immunoprecipitation. Asterisk (*) indicates a non-specific signal from antibodies (9D5). (C) Examples of capillary electrophoresis profiles for cytosolic RNA, P-TRAP RNA, and L10a-TRAP RNA from HEK293 cells. Two independent experiments were performed for each condition (#1, #2). The lower marker (LM) indicates internal standards (25 nt). RNA ladder (the Agilent RNA 6000 Pico kit) was used as a size marker.

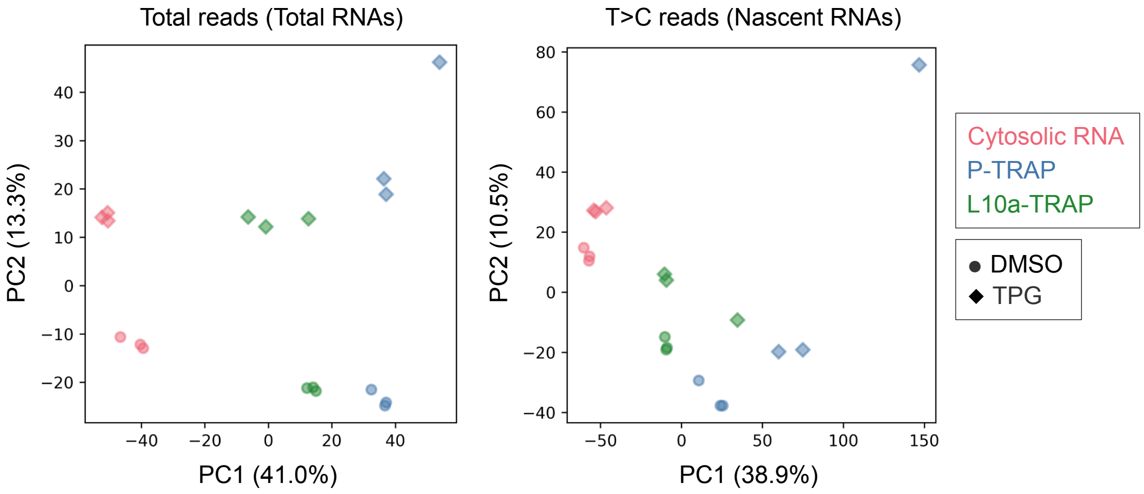


**Fig. S2.** Principle component analysis (PCA) of cytosolic RNA-seq (red), P-TRAP-seq (blue), and L10a-TRAP-seq (green) in the DMSO- (circle) and TPG-treated groups (diamond).

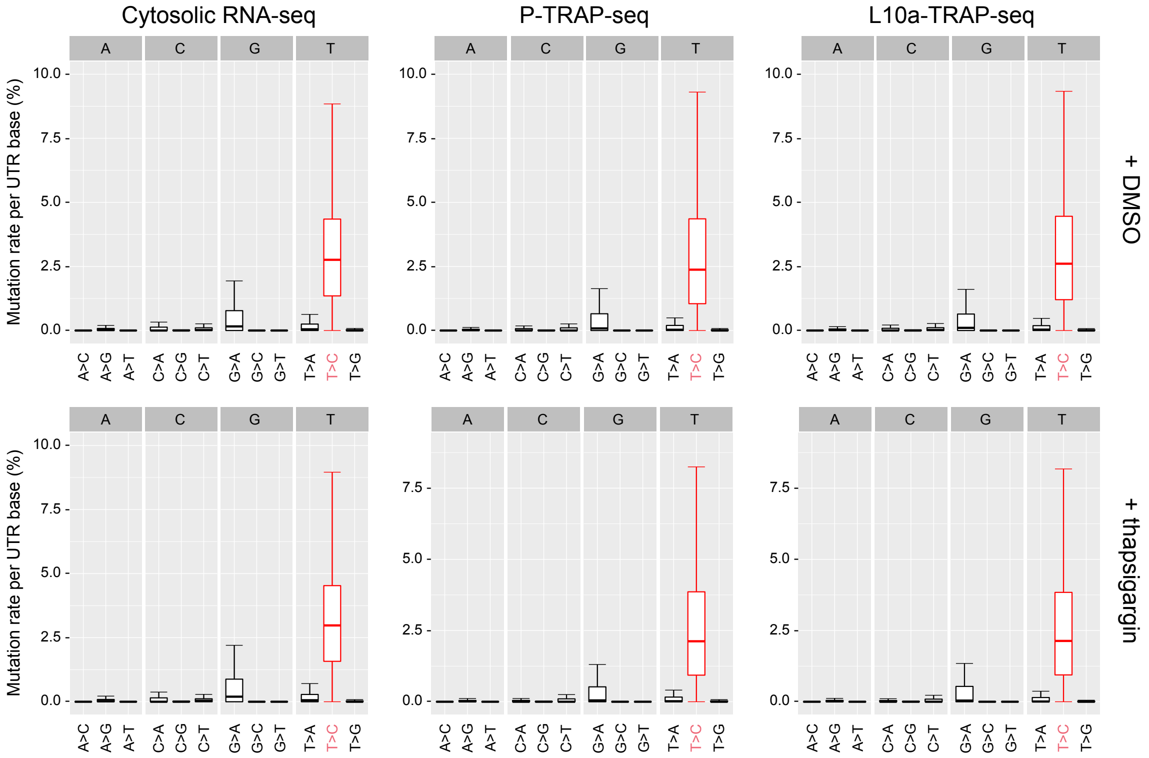


**Fig. S3.** Conversion rates in defined counting window-mapping reads of QuantSeq libraries, prepared from HEK293 cells after metabolic RNA labeling for 3 hours using 200 µM s4U.
